# Personalized electric field simulations of deformable large TMS coils based on automatic position and shape optimization

**DOI:** 10.1101/2024.12.27.629331

**Authors:** Torge Worbs, Bianka Rumi, Kristoffer H. Madsen, Axel Thielscher

**Affiliations:** Section for Magnetic Resonance, DTU Health Tech, Technical University of Denmark, Kgs Lyngby, Denmark; Danish Research Centre for Magnetic Resonance, Department of Radiology and Nuclear Medicine, Copenhagen University Hospital Amager and Hvidovre, Copenhagen, Denmark; Section for Cognitive Systems, Department of Applied Mathematics and Computer Science, Technical University of Denmark, Kgs Lyngby, Denmark; Sino-Danish College, University of Chinese Academy of Sciences, Beijing 100190, China

**Keywords:** TMS, electric field calculations, personalized dosing, Brainsway H-coils, MagVenture MST-twin coil

## Abstract

**Background:** Transcranial Magnetic Stimulation (TMS) therapies use both focal and unfocal coil designs. Unfocal designs often employ bendable windings and moveable parts, making realistic simulations of their electric fields in inter-individually varying head sizes and shapes challenging. This hampers comparisons of the various coil designs and prevents systematic evaluations of their dose-response relationships.

**Objective:** Introduce and validate a novel method for optimizing the position and shape of flexible coils taking individual head anatomies into account. Evaluate the impact of realistic modeling of flexible coils on the electric field simulated in the brain.

**Methods:** Accurate models of four coils (Brainsway H1, H4, H7; MagVenture MST-Twin) were derived from computed tomography data and mechanical measurements. A generic representation of coil deformations by concatenated linear transformations was introduced and validated. This served as basis for a principled approach to optimize the coil positions and shapes, and to optionally maximize the electric field strength in a region of interest (ROI).

**Results:** For all four coil models, the new method achieved configurations that followed the scalp anatomy while robustly preventing coil-scalp intersections on N=1100 head models. In contrast, setting only the coil center positions without shape deformation regularly led to physically impossible configurations. This also affected the electric field calculated in the cortex, with a median peak difference of ∼16%. In addition, the new method outperformed grid search-based optimization for maximizing the electric field of a standard figure 8 coil in a ROI with a comparable computational complexity.

**Conclusion:** Our approach alleviates practical hurdles that so far hampered accurate simulations of bendable coils. This enables systematic comparison of dose-response relationships across the various coil designs employed in therapy.

**Highlights:**

- automatic positioning and shape optimization of large deformable TMS coils
- ensures adherence to the head anatomy and prevents coil-head intersections
- enable automatic electric field maximization in target brain regions
- outperforms grid search for standard flat coils
- provides accurate computational models of four coils used in clinical practice

## Introduction

Transcranial Magnetic Stimulation (TMS) has been approved by several medical agencies as therapy against specific psychiatric diseases, including major depressive disorder and obsessive-compulsive disorder [1]. Interestingly, the clinically employed coil designs vary substantially and range from standard rigid figure-8 geometries to large and deformable coils [2–4]. Despite being used for the same clinical indications, the different designs induce electric fields in the brain that vary strongly in their spatial distribution and focality. The individual head and brain anatomies additionally influence the induced electric field (E-field) [5]. It is therefore important to understand how these two factors affect the therapeutic effects.

Personalized E-field simulations informed by structural magnetic resonance imaging (MRI) [5,6] can be useful tools to explore these questions, for example during clinical trials. However, this was so far practically difficult or infeasible for most of the therapeutically used large coils with complex shapes. So far accurate computational models of these coils for use in the simulations were mostly lacking. More importantly, the available simulation software does not have automatic means to prevent intersections between the coil model and the head model which will create physically impossible coil configurations and field distributions. Instead, time-consuming and practically tedious manual positioning of non-flat coil geometries on the head model is required. So far, shape adjustments of deformable coils were not supported, preventing realistic simulations of those coils.

The SimNIBS software package is an open-source tool for personalized E-field simulations [7] of transcranial magnetic and electric stimulation. While SimNIBS already includes many validated coil models [8], so far, it lacked support for coils with deformable and movable parts. In this paper, we introduce geometrically accurate models of the clinically used Brainsway H1, H4, and H7 coils and the MagVenture MST-twin coil [9]. We further describe a computationally efficient approximation of non-linear deformations of coil shapes by concatenated linear transformations. We use this to establish a principled optimization approach for fitting the coil position and shape to individual head shapes while avoiding intersections between the coil and the head models and, in case of the MST-twin coil, also between the two individually movable coil parts. The approach is generic and can be easily extended to further coil designs.

We assess two application scenarios, in which we evaluate the stability of our approach by reporting the average E-field distribution induced in the brain for 1100 head models. In the first scenario, the coil casing is fitted as close as possible to the head surface, to reach a physically feasible coil configuration close to an initial position provided by the user. In the second scenario, the coil position and shape are optimized to maximize the E-field in a cortical region of interest (ROI). For further validation, we apply the approach to a flat figure-8 coil and compare the optimized E-field with the results of a standard grid search.

## Methods

### Modeling of the Brainsway H1, H4 and H7 coils and the MagVenture MST-twin coil

Physical samples of the Brainsway H1, H4 and H7 coil models were scanned in a clinical computerized tomography (CT) scanner, whereby the coils were carefully placed to avoid deformations. The wire paths were then traced manually in the scans (Suppl. Material A). Calculations of the magnetic fields and magnetic vector potentials of the coils were then implemented based on numerical solutions of line integrals [10] using the fast multipole method for improved efficiency [11]. A wire was approximated by a single line element placed in its center with a resolution of two line elements per millimeter. This resulted in a numerical error below 0.04% compared to reference cases with very dense sampling. This choice enabled good numerical accuracy (Suppl. Material A) while maintaining computational efficiency. When positioning the H1, H4 and H7 coils physically on heads, the resulting deformations occurred as bends of specific wire paths, which could be well approximated in the models by rotations around “bending axes” or, more generally, affine transformations. If needed, more complex deformations were represented by chaining affine transformations that are successively applied to different sub-parts of the wire paths. By this procedure, coil deformations can be efficiently expressed as a small number of affine transformations. Based on pilot tests on various head shapes, we visually identified the deforming wire paths and defined appropriate bending axes and physically feasible transformation ranges. The coil wires are attached to a fabric cap and include soft padding. After fitting the coils firmly on a head using the integrated straps, we measured the minimal gap between skin and wires caused by the cap and padding materials. For modeling this gap, we then created surfaces around the wire paths (Fig. 1A) to account for the padding when computationally fitting the coil model on head models. We assessed the accuracy of the modeled deformations and gaps by performing a CT scan of the H1 coil fitted on a ball with a 200 mm diameter, confirming that the computational coil model approximated the physically occurring deformation well (Suppl. Material B).

**Figure 1:**
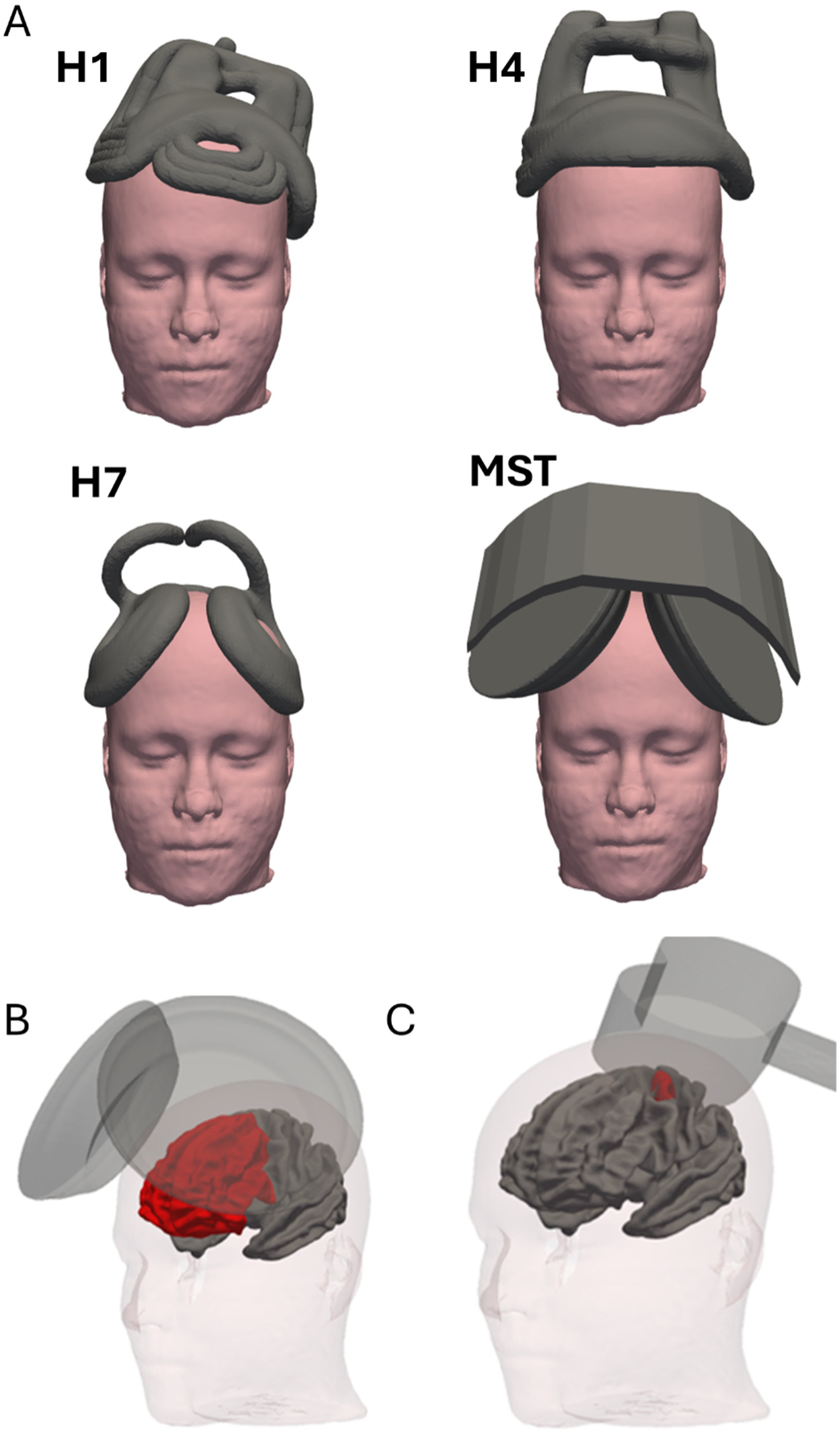
A) Initial placements on the MNI head for the distance optimization of the Brainsway H1, H4 and H7 coils, and MagVenture MST-Twin coil. B) Initial coil placements on the MNI head for optimization of the average electric field magnitude (abbr.: mean(|E|) within a bilateral prefrontal ROI (shown in red) for MagVenture MST-Twin coil. C) Left precentral ROI covering the handknob for use with the MagVenture Cool-B65 coil. The masks are defined on the "fsaverage" cortical surface and transformed to the subject space via surface-based registration [34].

The MagVenture MST-twin coil has two “sub-coils” that are connected to a guide rail, which determines the range of feasible positions of the two sub-coils. Using an existing computational model of an MST sub-coil [8] and measurements on a physical coil sample, we created a model of the complete MST coil consisting of representations of the two sub-coils and the guide rail (Fig. 1A). The feasible positions of the sub-coils were tested on the physical coil sample and represented by defining suited linear transformations and parameter ranges.

A new Json-based file format (“.tcd” – TMS coil definition, Suppl. Material C) was created for the generic representation of the four coil models in the SimNIBS simulation environment, also substantially simplifying addition of further coil models with complex shapes.

### Optimization of coil position and shape: Cost functions

We implemented automatic optimizations of the coil position and shape, supporting two different application scenarios. In the first scenario, the objective is to smoothly fit the coil casing on the head surface, starting from an initial position provided by the user and then adjusting the position and deformation of the coil. Thereby, a minimal distance to the head surface is ensured and self-intersections (e.g. intersections of the two sub-coils of the MST-twin coil) are avoided. This was formalized in a cost function

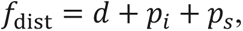

where the distance *d* is determined as the average distance of predefined positions on the relevant coil wires or casing parts that are close to the head (Suppl. Fig. S4). The intersection penalty variable *p*_s_ is based on the calculation of the intersection volume between the coil casing volume and the head volume, whereby deeply intersecting parts are weighted more to support faster convergence of the optimization. The self-intersection penalty variable *p*_s_ is the total intersection volume between sub-parts of a coil.

The second scenario aims at the maximization of the electric field strength in a ROI in the brain while preventing intersections between the coil and head and coil self-intersections. The corresponding cost function has the form:

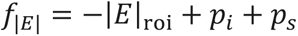

The variable |*E*|_roi_ is the average electric field strength in the ROI. Appropriate weighting constants for the different parts of the cost functions were set in pilot tests to ensure robust convergence of the optimization and avoidance of intersections. Additional information about the definition and implementation of the different parts of the cost functions is stated below and in Supplementary Material D.

### Optimization framework

The optimization was performed with a combination of global optimization (DIRECT or the faster locally-biased DIRECT-L [12]) followed by additional local optimization using a quasi-Newton method (L-BFGS-B [13]). Unless stated otherwise, DIRECT-L and L-BFGS-B were chosen as the default methods for the results reported in the main paper, which provided a good tradeoff between the number of function evaluations and the optimization result. The default method was compared to DIRECT and L-BFGS-B as reference.

Parameter bounds were set according to the physically feasible coil deformation ranges. In addition, the coil center positions and orientations were restricted to reasonable ranges, as determined in pilot tests. The ranges for coil translations, rotations, and deformations are listed in Supplementary Table S4.

To evaluate the general performance of our approach, we compared our e-field optimization results for the non-flexible MagVenture Cool-B65 coil to a standard grid search approach [14]. As local optimization turned out to be sufficient to ensure adequate optimization, the MagVenture Cool-B65 results reported in the main paper only utilized local optimization with L-BFGS-B, while further results are shown in Supplementary Material E.

### Selection of initial coil positions and orientations, and definition of target ROIs

The initial coil placements for optimization were set according to the coil model and optimization goal. Unless stated otherwise, the coil positions and orientations were manually defined in MNI space (Fig. 1A), in an effort to minimize the coil-head intersections, and then non-linearly transformed to the individual subject spaces using SimNIBS functions. These coil settings are included as the “baseline” in the results to illustrate how well non-deformable coil models optimized once on a template would work.

- *Distance* optimization of the Brainsway H1 coil: The initial position of the coil center (see Suppl. Fig. S5 for details) was chosen to be above the center of area BA46 of the MNI template head (MNI coordinates [-44, 40, 29]), a common target in the treatment of MDD [15]. The orientation of the coil was manually adjusted along the left-right and anterior-posterior axes to minimize the intersection of the coil with the MNI template.
- *Distance* optimization of the Brainsway H4 coil: The initial coil position was selected to mimic the clinical guidelines for coil placement as provided by Brainsway. The coil center was placed on the MNI template head above the group-average activation site of the FDI muscle (MNI coordinates [−41, −7, 63] [16]), then moved medially to a position above the interhemispheric cleft and finally projected 6 cm anteriorly, resulting in the MNI position [0, 53, 63] (Suppl. Fig. S6). The orientation of the coil along the left-right axis was manually optimized to minimize coil-head intersections. The rotations along the other two axes were chosen to ensure symmetric placement of the coil above both sides of the head.
- *Distance* optimization of the Brainsway H7 coil: A position between the medial parts of the primary motor areas that contain the leg representations was visually determined and the closest scalp position was chosen (MNI coordinates [-5, -20, 99]). The treatment position was then found by moving 4 cm in the anterior direction (MNI coordinates [-5, 20, 87]; Suppl. Fig. S7), according to clinical guidelines of the company.
- *Distance* and *E-field* optimization of the MagVenture MST-Twin coil: Clinical applications aim to place the two coil halves above the F3/F4 electrode positions, respectively [17,18]. The coil center was thus placed at the Fz electrode position between F3/F4 (Suppl. Fig. S8), and the coil model was additionally moved in inferior-superior direction to achieve a placement of the two sub-coils approximately on the skin surface. As the MST-Twin coil is designed for seizure induction by stimulating frontal areas, a large bi-lateral frontal ROI was used as target for the *E-field* optimizations (Fig. 1B).
- *E-field* optimization of the MagVenture Cool-B65 coil: As the Cool-B65 coil is used for focal stimulation, we defined the hand knob area of the left precentral gyrus as an example target ROI (Fig. 1C). The coil position was directly initialized in subject space by automatically determining the skin position closest to the brain ROI (termed “auto-init” in the following). Alternatively, the coil position was centered over the hand knob (MNI coordinate [-32, -26, 59]), and orthogonally projected to a 4 mm skin distance from the scalp after transformation to subject space. The coil handle was oriented backwards and approximately 45 degrees lateral from the mid-line thereby ensuring that it pointed in the direction opposite to EEG 10-10 position FCz. The results for the “auto-init” option are reported in the main part of the paper, while the subject-specific settings serve as comparison and are reported in Supplementary Figure S10C and Supplementary Table S2.

### Software Implementation

The methods and coil models presented in this paper will be published as open source software in the next SimNIBS release. Also examples on how to define additional custom coil models will be included. The intersection tests are based on voxel masks (1 mm³ isotropic) of the coil and head surfaces that are automatically voxelized during runtime [19]. Intersection volumes between two masks are then efficiently determined by interpolating one mask to the voxel space of the second mask (using the map_coordinates function in SciPy 1.13.1 [20]), according to the linear transformation representing the shape deformation, and summing overlapping voxels. The evaluation of the *distance* term was implemented in a similar way, but weighted by the intersection distance (Suppl. Material D). The speed of the FEM calculations of the *E-field* in the brain ROIs was optimized for computational efficiency using a similar approach as outlined in Cao, Madsen et al [21]. The implementations of DIRECT, DIRECT-L and L-BFGS-B in SciPy 1.13.1 [20] were used. Visualizations were created with Matplotlib 3.9.2 [22] and PyVista 0.44.1 [23].

### Evaluations of the Optimization Performance

To evaluate the stability of the optimization approach for varying head shapes, results were assessed on the Human Connectome Project database of young healthy adults (N=1100) [24]. SimNIBS *charm* [25] was used for the creation of the head models from the T1- and T2-weighted structural MRI images, incorporating reconstructions of the pial and white matter surfaces from FreeSurfer [26] for more accurate representations of narrow sulci in the head models. The resulting head meshes consisted of ∼4.6 million tetrahedral elements representing seven tissue types (white matter, gray matter, cerebrospinal fluid, compact bone, spongy bone, scalp, and eye balls), and their default conductivity values were used in the simulations (0.126 S/m, 0.275 S/m, 1.654 S/m, 0.008 S/m, 0.025 S/m, 0.465 S/m, and 0.5 S/m). All simulations were performed for a rate of change of the coil current of 1 A/µs, and the E-field strength was evaluated on the central gray matter surfaces of two hemispheres halfway between the pial and white matter surfaces. All head meshes were visually inspected for quality assurance. In addition to the results obtained on the large dataset, speed tests were performed on a desktop computer (Ubuntu 22.04, Intel i7-11700, 16 cores, 32GB RAM) using the public SimNIBS *ernie* dataset with standard conductivities to demonstrate expected run times in practical settings. The target ROIs for the *E-field* optimizations were defined in the fsaverage surface space and automatically transformed to the individual subject spaces using SimNIBS functionality.

The evaluations were based on the following criteria:

- *Distance optimizations:* The effect of the optimization on coil position and shape was indexed using the *d* metric (distance, as defined above) before and after optimization. In addition, a signed distance measure was calculated between the head mesh and the coil casing, whereby negative values indicated the maximal (or deepest) intersections of the coil with the head, while positive values correspond to the minimal distance between the coil casing and the scalp in case of no intersections. For evaluating the effect of the optimization on the calculated electric fields, simulations were performed for the baseline configuration and after optimization. The electric field strengths on the central gray matter surfaces of both hemispheres (i.e. on surfaces midway between the pial surface and gray-white matter boundary) were determined and the differences between baseline and the optimized coil configuration were calculated and scaled relative to the 99.9-percentile of the electric fields after optimization. The results were mapped to the fsaverage surface space to obtain group difference maps. The computational efficiency of the optimization was determined using the average (+/- standard deviation) number of function evaluations. In addition, the runtime on the test desktop computer for the SimNIBS *ernie* dataset was evaluated.
- *E-field optimizations of the MagVenture MST-Twin coil:* The average electric field strength in the ROI (Fig. 2a) at baseline and after optimization was compared. In addition, signed distance measures were calculated for both cases.
- *E-field optimizations of the MagVenture Cool-B65 coil:* Signed distance measures were calculated to confirm the absence of intersections after optimization. In addition, we compared the average electric field strength in the ROI achieved with the new optimization to the results of standard grid search implemented in SimNIBS [14]. The latter was feasible for this coil as the Cool-B65 is flat and does not deform. Grid search was performed on the full dataset (N=1100) with a grid spacing of 5 mm and 12 orientations per position with an angular spacing of 15° (585 simulations per grid). In addition, a higher-resolution grid search was performed on a random subset of N=48 subjects with a spacing of 2 mm and 36 orientations per position with an angular spacing of 5° (11285 simulations per grid). The grid search only optimizes the position of the coil center on the skin surface and the orientation of the coil handle, while the coil is always selected to be parallel to skin surface under the coil center to simplify the search. However, this resulted in skin intersections for several of the grid search results due to irregular head shapes. To enable a fair comparison, the final coil position was moved orthogonally away from head in these cases until the intersections were resolved, and the electric field in the ROI was reevaluated.

**Figure 2:**
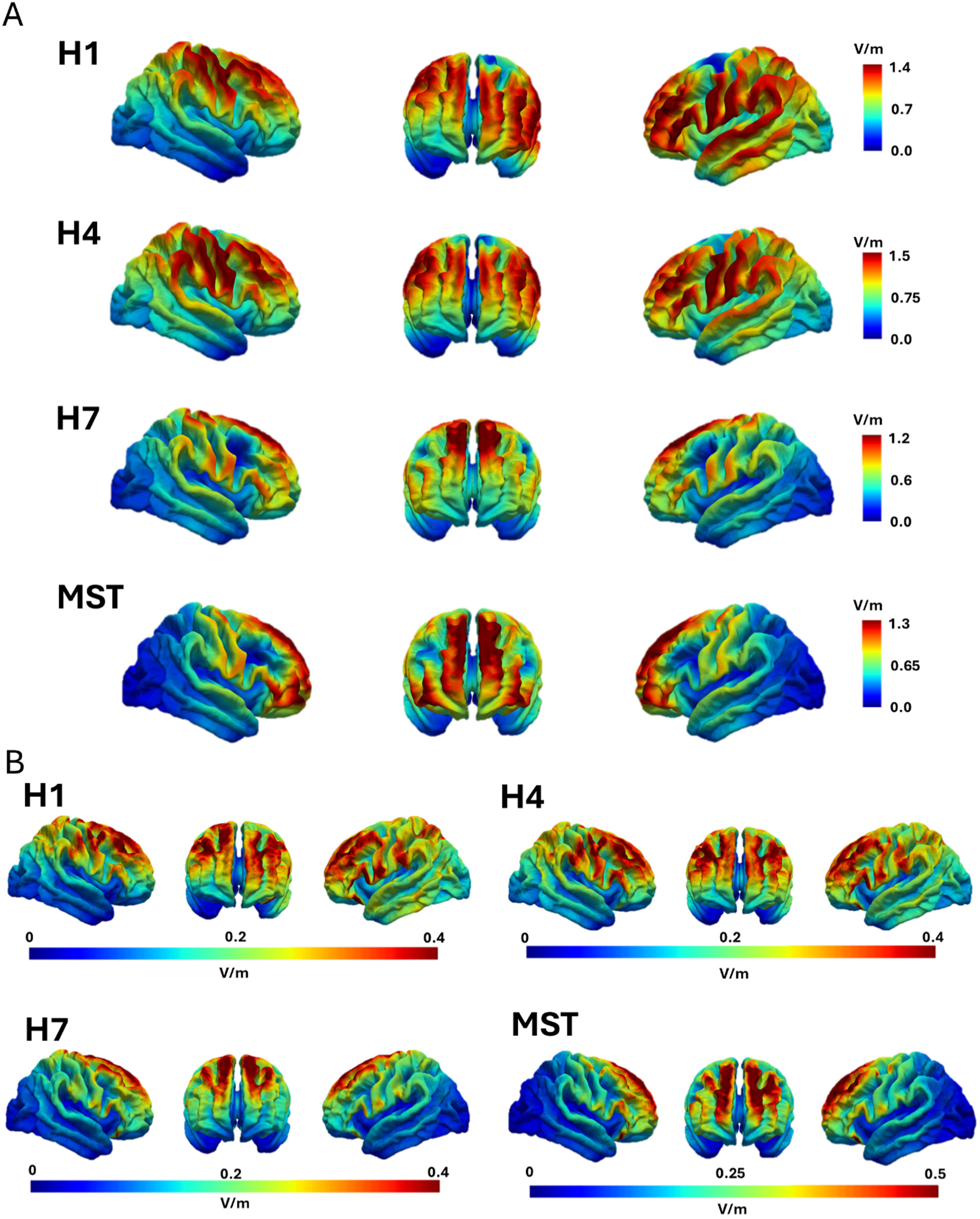
E-fields of the H1, H4, H7 and MST-Twin coils after coil distance optimizations. A) Median across N=1100 head models of the induced electric field strength. B) IQR across the 1100 head models.

## Results

### Distance optimization

The median electric field distributions across the 1100 head models and the interquartile ranges (IQR) after distance optimization are shown in Figure 2 for the H1, H4, H7 and MST coils. Table 1 lists the peak E-field strengths in the cortex (defined as the 99.9%ile), focality measures (defined as the gray matter volume where the E-field strength exceeds 50% of the peak strength) and depth measures (defined as the radial distance from the brain surface to the deepest point in gray matter where the E-field strength is half of the peak strength) [27,28].

**Table 1.** Fields summary of the optimization results for distance optimization (H1, H4, H7, MST-Twin) and electric field optimization (MST-Twin and Cool-B65). The values are shown as the medians ± the interquartile range across the subject pool.

|  | Distance optimization |  |  |  | E-field optimization |  |
| --- | --- | --- | --- | --- | --- | --- |
|  | H1<br>DIRECT-L+<br>L-BFGS-B | H4<br>DIRECT-L+<br>L-BFGS-B | H7<br>DIRECT-L+<br>L-BFGS-B | MST-Twin<br>DIRECT-L+<br>L-BFGS-B | MST-Twin<br>DIRECT-L+<br>L-BFGS-B | Cool-B65<br>L-BFGS-B |
| **Peak \|E\| (V/m)** | 1.9 ± 0.1 | 2.2 ± 0.2 | 1.8 ± 0.2 | 2.1 ± 0.2 | 2.1 ± 0.2 | 1.14 ± 0.2 |
| **Focality (cm<sup>3</sup>)** | 108.0 ± 18.1 | 89.0 ± 16.6 | 37.8 ± 9.0 | 41.5 ± 8.3 | 43.1 ± 8.1 | 12.8 ± 3.8 |
| **Depth (mm)** | 22.6 ± 2.1 | 21.5 ± 1.9 | 22.9 ± 2.4 | 21.5 ± 2.0 | 21.3 ± 2.0 | 16.7 ± 1.6 |

The optimization reduced the interindividual spread of the coil-scalp distances compared to the initial positions (Fig. 3A), suggesting that it led to a more consistent fit of the coil models on the head surfaces. It successfully removed all intersections of the coils with the heads and achieved consistent minimal distances to the skin surfaces (Fig. 3B). The results shown for the initial positions demonstrate that using coil positions transformed from a group template space and ignoring deformations regularly results in impossible positions on the individual heads. The median of the differences between the E-field strengths induced in the cortex for the optimized vs initial positions reach 12% (MST coil) and 16% (H1, H4 & H7 coils) of the maximal E-field strength (Fig. 3C). Of note, strong differences occur also in brain regions that are implicated in the therapeutic effects, such as the prefrontal cortex for the H1 and H4 coils. Looking at the 90-percentile (Suppl. Fig. S9) reveals differences exceeding 20% for all coils in 110 of the 1100 head models. Overall, these results suggest that distance optimization is important to reach plausible coil configurations and that this has clear effects on the E-field distribution on the cortex.

**Figure 3:**
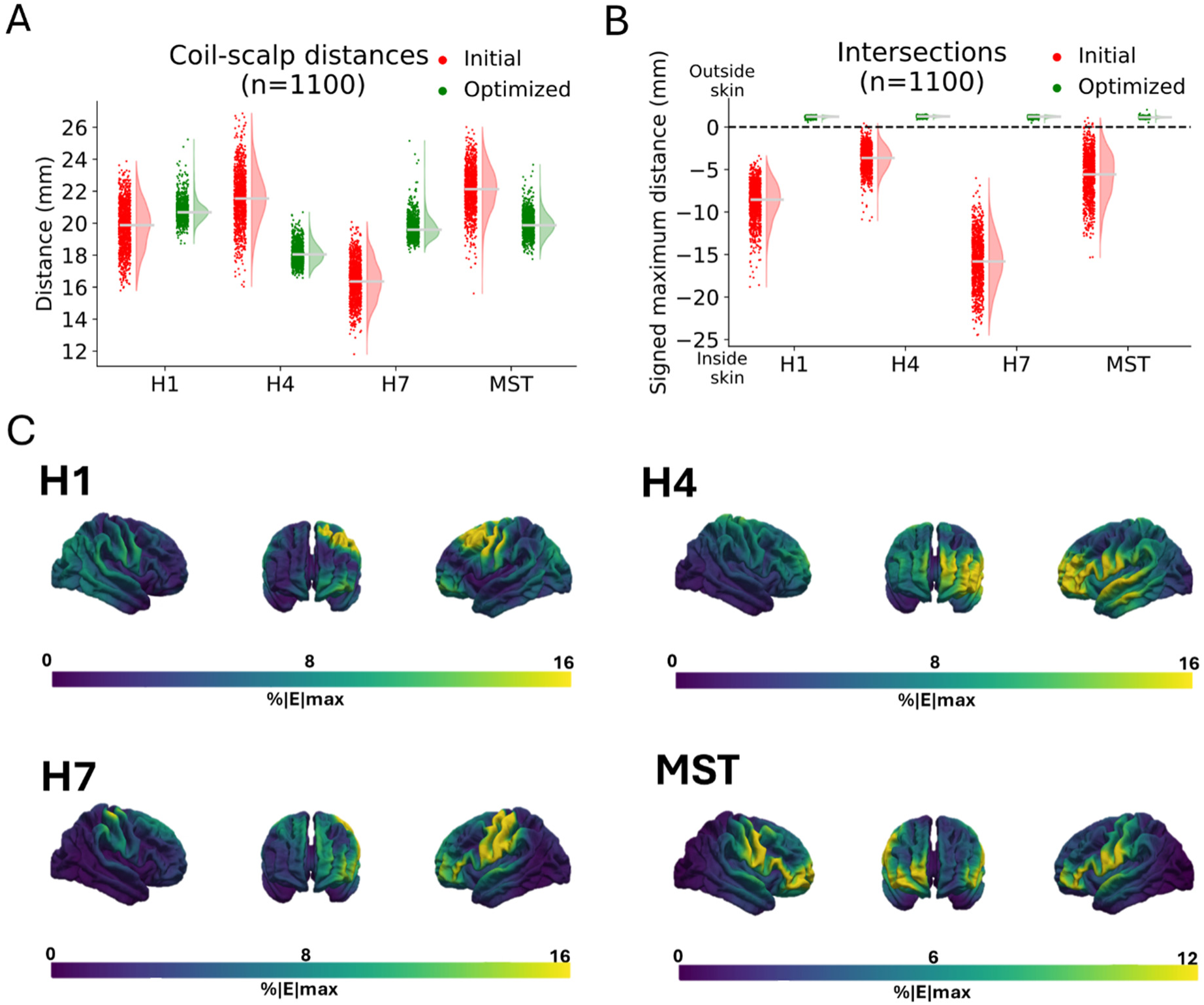
A) Coil-scalp distances before and after optimization. The distance is calculated as the mean distance between the pre-defined sets of positions on the coil casings and their nearest scalp position for each subject. B) Maximally occurring intersections of the coil with the head before and after distance optimization. Signed distances are reported, with negative values indicating intersections, and positive values the minimal distance between coil casing and scalp in case of no intersections. Specifically, for each individual optimization result, the minimal value of the signed distances between the scalp and any of the predefined positions on the coil casing is shown. This corresponds to the deepest intersection or, in case no intersection occurred, positive values indicate the minimal distance between coil casing and scalp. C) Median of the relative differences between the electric field strengths for the optimized and initial coil settings.

Comparison of the employed optimization method (DIRECT_L&L-BFGS-B) with a more extensive optimization (DIRECT with stricter convergence settings, followed by L-BFGS-B) reveals mostly equal performance, with the differences in the distance cost function being centered around zero (Suppl. Fig. S10A). The more extensive optimization reaches cost function values that are better by 10% or more only in a few outlier cases. On the other hand, DIRECT_L&L-BFGS-B required on average far less function evaluations compared to DIRECT&L-BFGS-B (e.g., 2137 vs 7816 for the H1 coil; see Suppl. Table S1 for all results). The total time for the distance optimizations on the desktop computer stayed below 3 minutes for all four coil models and the memory requirements stayed below 4 GB (DIRECT_L&L-BFGS-B and *ernie* head model; Suppl. Table S3).

### Electric field optimization

The median electric field distributions and their IQR maps for the MST-Twin coil after optimization are shown in Figure 4A&B. Table 1 lists the corresponding peak electric fields and focality results. Figures 4E&F show that the optimization led to an increase of the average E-field strength in the bilateral prefrontal ROI from ∼0.72 V/m (median across subjects) to ∼0.75 V/m while robustly avoiding coil-scalp intersections. The median of the differences between the E-field strengths in the cortex reaches 14% of the peak E-field strength (Fig. 4C). The medians between E-field strength from the position optimization and electric field optimizations differ with up to ±0.1 V/m across subjects (Fig. 4D).

**Figure 4:**
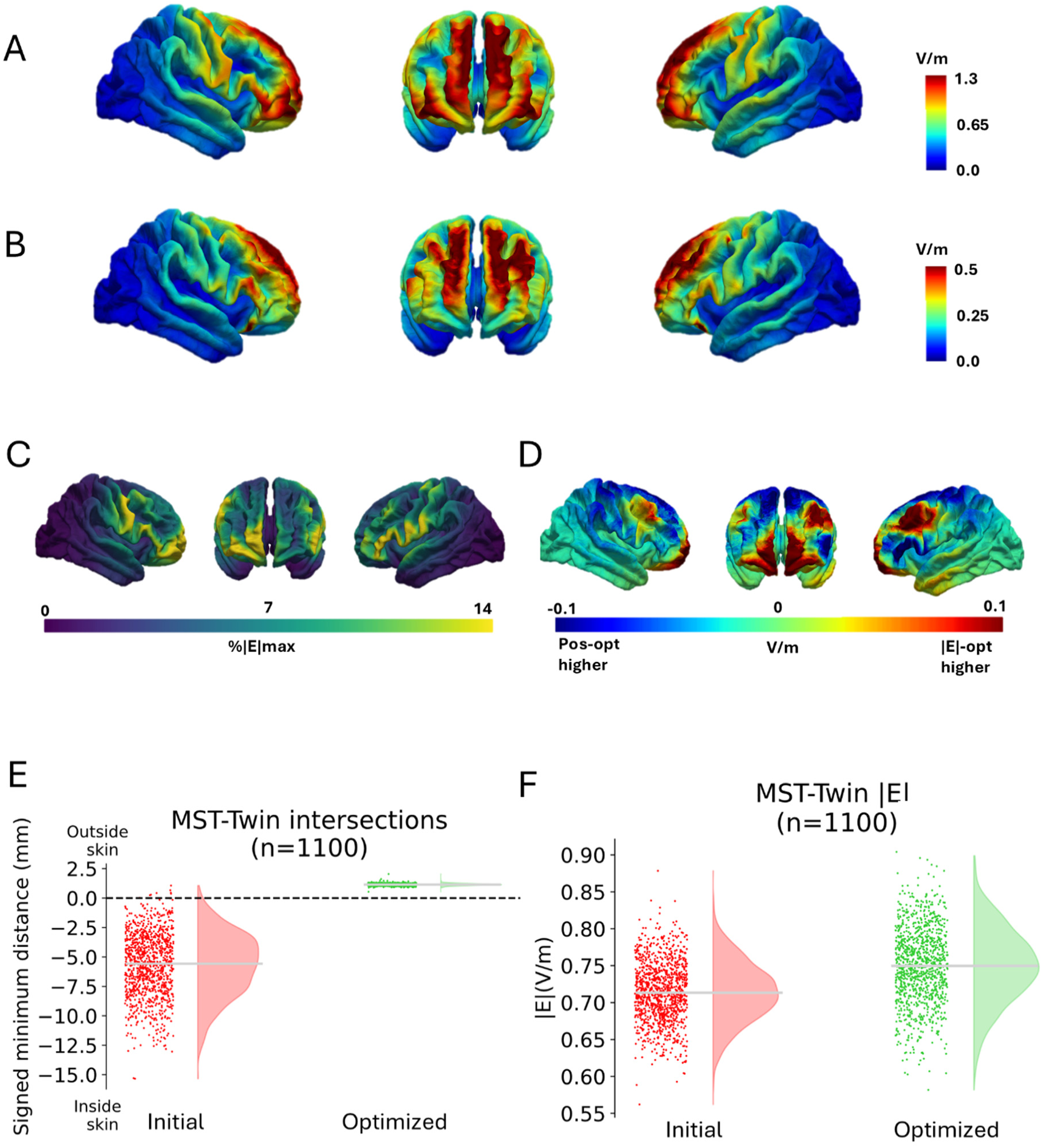
Results of the optimization of the mean electric field strength in the bilateral prefrontal ROI for the MST-Twin coil. A) Median across N=1100 head models of the electric field strength after optimization. B) IQR across 1100 head models. C) Median of the relative differences between the electric field strengths for the optimized and initial coil settings. D) Average difference in electric field strength between electric field strength-optimized and position-optimized MST-Twin coils (Figure 4A and 3A) E) Mean electric field strengths in the ROI before and after optimization. F) Maximal intersections of the coil with the head before and after optimization. Signed distances are reported, with negative values indicating the maximal depth of the intersection, and positive values the minimal distance between coil casing and scalp in case of no intersections.

Comparison of the DIRECT_L&L-BFGS-B optimization with the more extensive DIRECT&L-BFGS-B optimization again reveals similar performance, with the differences in the achieved costs centered around zero and only a few cases where the DIRECT&L-BFGS-B optimization is better by 10% or more (Suppl. Fig. S10B). However, DIRECT_L&L-BFGS-B required on average substantially less function evaluations (2508 vs 4921; Suppl. Table S1). The total time for the E-field optimization with the *ernie* head model was around 30 minutes on the desktop computer and the memory requirements were below 8 GB (Suppl. Table S3).

The E-field strength induced by a Magventure Cool-B65 coil in a left precentral gyrus ROI covering the primary motor hand area (handknob) was additionally optimized using only the local L-BFGS-B search and compared to the results of naïve grid searches. Figure 5A shows the differences of the achieved E-field strength in the ROI when compared to a lower-resolution grid search for all 1100 head models. Figure 5B shows the corresponding results compared to a high-resolution grid search for a sub-group of 48 head models. Our optimization approach reliably prevented coil-scalp intersections (data not shown) and generally performed slightly better than the grid search. This can be explained by the fact that the grid search enforces a tangential coil orientation relative to the skin surface under the coil center while our approach optimizes all 6 degrees of freedom of the coil position.

**Figure 5:**
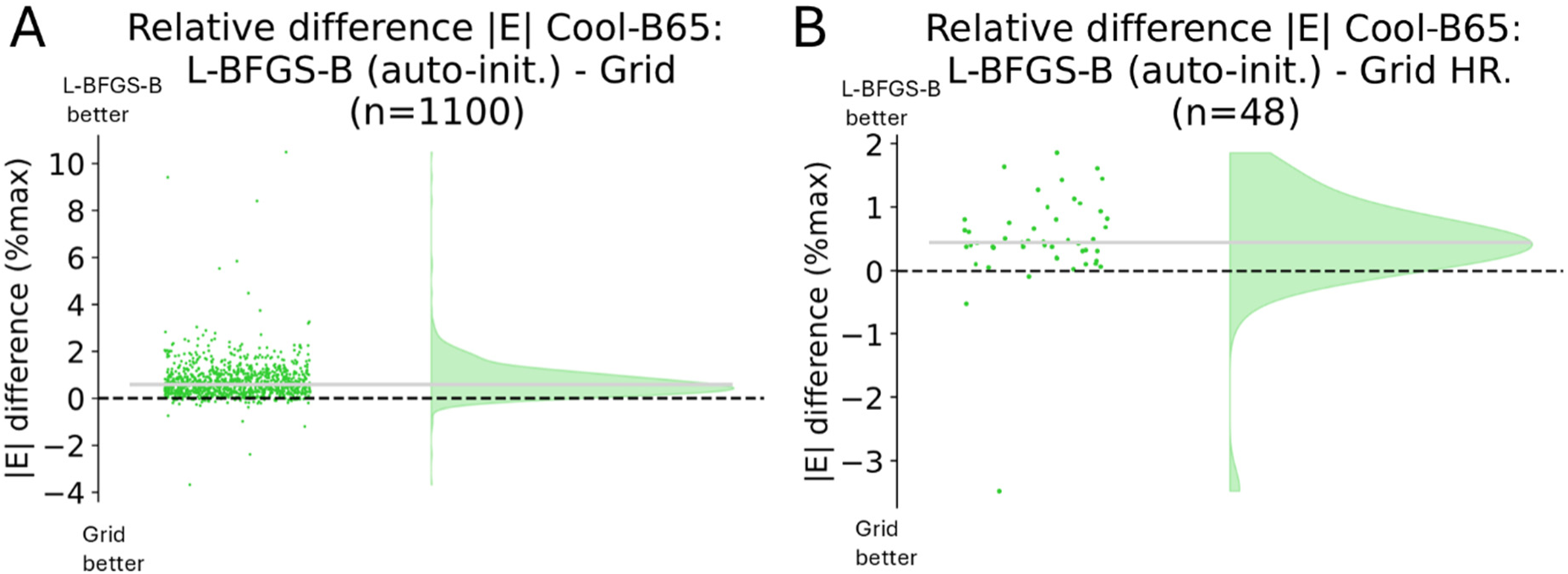
Results of the optimization of the mean electric field strength in the handknob ROI for the MagVenture Cool-B65 coil. The differences of the mean electric field magnitudes in the ROI obtained by our optimization approach versus a grid search are shown. A) Comparison to results of a “coarse” grid search and N=1100 subjects. B) Comparison to results of a higher-resolution grid search (N=48).

Using only local L-BFGS-B search was generally sufficient to reach stable optimization results for the Cool-B65 coil, which did on average not improve further for more extensive search strategies (Suppl. Fig. S10C). Using the most extensive search with DIRECT&L-BFGS-B seemed to slightly reduce the already very low number of outliers where grid search was better by 1% or more. Optimization with L-BFGS-B and automated initial position required on average 423 function evaluations compared to the low- and high-resolution grid searches with 585 and 11285 evaluations (Suppl. Table S2). The total time for the E-field optimization for the Cool-B65 coil and the *ernie* head model stayed below 8 minutes on the desktop computer and the memory requirements were below 8.5 GB (Suppl. Table S3). Overall, the results for the Magventure Cool-B65 coil suggest that the new optimization approach can serve as a general approach for all coil types including standard flat and rigid coils.

## Discussion

### Summary of findings

We introduced the first method to automatically optimize the shape of deformable TMS coils and validated its performance on a large dataset of 1100 head models and four new models of large TMS coils that are used in approved therapies or clinical trials. We demonstrated that it robustly avoided coil-scalp intersections while achieving coil configurations that fitted closely to the head surface or maximized the E-field strength in a target ROI. The reported group median and IQR maps for H1, H4, H7 and MST coils will be provided as online resources.

In contrast, we showed that the use of coil positions that were determined via transformations from a group template space and did not include deformations regularly resulted in physically impossible positions on the individual heads. This resulted in notable differences in the E-field distributions between the two cases that also occurred in potential therapeutically relevant brain areas. The number of function evaluations and total runtime on a normal desktop computer were low enough to allow for standard practical use.

We additionally compared our approach to a previously published grid search approach, with the aim to maximize the E-field strength induced by a flat and rigid coil (Magventure Cool-B65) in a small target ROI. Our approach performed at least as good as the grid search and required fewer function evaluations on average. This suggests that the new optimization method can serve as a general-purpose approach for all coil types.

### Comparison to published work

We compared the performance of our method to standard grid search. Alternative optimization approaches are the auxiliary dipole method (ADM, [29]) that can maximize the average E-field in a ROI and its recent extension [30] which determines the full E-field distribution in the ROI. In principle, the FEM-based E-field calculations in our approach could be replaced by these methods. As both methods use precalculations, it would depend on the number of function evaluations whether this would result in a speedup of the overall time required for optimization. More importantly, the methods are restricted to E-field calculations and do not allow for coil-scalp intersection testing or the handling of deformable coils.

The electric field distributions of the Brainsway H1, H4&H7 coils and the MagVenture MST twin coils were characterized by a few prior publications [27,31–33] using a variety of approaches such as measurements in simplified saline-filled phantoms [33], simulations using a spherical multi-layer model to coarsely mimic the head anatomy [27] or a low number of anatomically detailed head models [31,32]. The coils were modelled based on geometric information provided by the manufacturers and fitted manually to the skin surfaces. Our study complements these findings by ensuring the accuracy of the coil models, systematically optimizing the fit of the coils on the skin surfaces and assessing the E-field characteristics in a large sample of anatomically detailed head models. Differences in the applied metrics make a direct comparison with these prior findings difficult. However, our focality and depth estimates for the H1 coil were found to be comparable to a previous approximation based on a spherical head model [27]. Another study using two anatomically detailed head models reports notably larger depth estimates, but also included the electric field in white matter in the estimations [32].

### Limitations

To the best of our knowledge, the presented method is the so far only automatic approach to realistically place deformable coil models and to systematically avoid coil-head intersections. Our approach well approximated the true shape of an H1 coil put on a ball, which led to stronger coil deformations than when placing it on heads in our pilot tests. The H4 generally deformed less in our tests. The H7 and MST-Twin coils consist of two sub-coils that do not or hardly deform but only change their relative positions, which could be accurately modelled by linear transformations. We thus suggest that the tested H1 deformations serve as a reasonable worst-case scenario and are confident that our modeling approach captures practically feasible coil configurations with good accuracy. However, coil deformations in practice also depend on, e.g. how strongly the straps of the H-coils are tightened, so that the exact correspondence between the modelled and real coil configurations would need additional experimental assessment.

Our approach allows for flexibility in terms of the optimization methods utilized and their parameter settings. The results in the main paper are based on settings that balance accuracy and computational efficiency. More exhaustive search strategies gave little additional gain, except for slightly reducing the already very low numbers of outlier cases in which the balanced settings performed worse to a notable extent. However, we recommend choosing the optimization settings and by that the tradeoff between runtime and avoidance of putative outliers with the specific research question in mind.

The reported electric field distributions might deviate from those achieved in clinical practice due to differences in coil positioning strategies. We place the coil centers at average positions defined in MNI space that might lead to deviations compared to positions that are defined using features of the individual brain or head anatomy. Specifically, for the H1 coil, we chose its center to be above area BA46 in MNI space, while it is placed relative to the motor hot spot of the finger muscles in practice.

### Future objectives

We presented a new framework for the simulation of large and deformable TMS coils that systematically prevents coil-scalp intersections. We tested it with accurate computational models of four therapeutically employed coils and a dataset of 1100 head models. We suggest that this will facilitate evaluations of the dose–response relationships for these coils, including estimations of interindividual differences of the induced E-fields and systematic comparisons of induced E-fields across different coil models. Additionally, our approach can be used for the principled optimization of coil positions and configurations to maximize the stimulation of a target brain area. The method and coil models will be provided as part of SimNIBS together with comprehensive example scripts demonstrating their usage. Moreover, examples for creating custom computational models of rigid and deformable coils will be included.

## Acknowledgements

We thank BrainsWay Ltd (Jerusalem, Israel) for providing physical samples of the H1, H4 and H7 coils and MagVenture A/S (Farum, Denmark) for providing a sample of the MST-Twin coil.

## Funding

AT was supported by Innovation Fund Denmark (Grand Solutions grant 9068-00025B “Precision-BCT”) and the Lundbeck Foundation (grants R313-2019-622 and R244-2017-196). TW and AT received support by the National Institute of Health (grant R01MH128422).

## Supplementary Material A: Coil implementation based on CTs

Physical samples of the Brainsway coils (H1, H4, and H7) were provided by the manufacturer. These coils underwent CT scanning on a Siemens SOMATOM Definition Flash using a sequence with 0.5×0.5×3 mm resolution, 140 keV, 600 mAs and metal artifact reduction. The resulting CT scan data was converted into NIfTI format for further processing using custom software written in Python 3.8. The origins and axes directions of the NIfTI images were set manually according to the information about the coil centers provided by the manufacturer.

For each coil, a binary mask was generated by setting an appropriate intensity threshold to isolate the metal wires and transformed into triangle surfaces using a marching cube algorithm. The resulting surfaces represented the circumferences of the coil wires, and the central paths of the wires were obtained by shrinking the surfaces along their local normal directions. These central wire paths were manually tracked, yielding precise wire path coordinates. Subsequently, these wire path coordinates were interpolated using b-splines to obtain a desired resolution (2 elements/mm). The coil center position and coil coordinate system were defined based on manually identified coil landmarks in the CTs (mounting screws, brackets on the coil, see Fig. S4 below). The interpolated and aligned data points were transformed into deformable coil models, in line with the coil generation examples included in SimNIBS 4.5. The “casing thickness” to account for the fabric cap and paddings and the coil deformations were defined as outlined in the main paper.

To estimate the induced error from representing the coil wire by a spatially discretized line current along the central wire path, two convergence tests were performed with a round coil (radius of 50 mm; wire diameter of 5 mm) placed above electrode position Cz of the standard SimNIBS *ernie* head model (Suppl. Figure S1). The errors of the electric field magnitude and the electric field vectors were calculated in relation to reference cases. The first test assessed the error due to modeling the wire by a single line current along the central wire path. For that, the wire was modelled by an increasing number of line currents placed in spherical layers (Fig. S1A). The resolution along the wire paths was kept constant at 3.2 elements/mm. The reference consisted of 8 layers with 169 lines. Using one central line already resulted in a very low error that is practically not relevant (≈0.038% error).

The second test assessed the convergence when increasing the resolution along the wire path, using 64 elements per mm as reference. Two elements per mm resulted in an error of ≈0.001% and were selected when modelling the wire paths of the H coils.

**Supplementary Figure S1.**
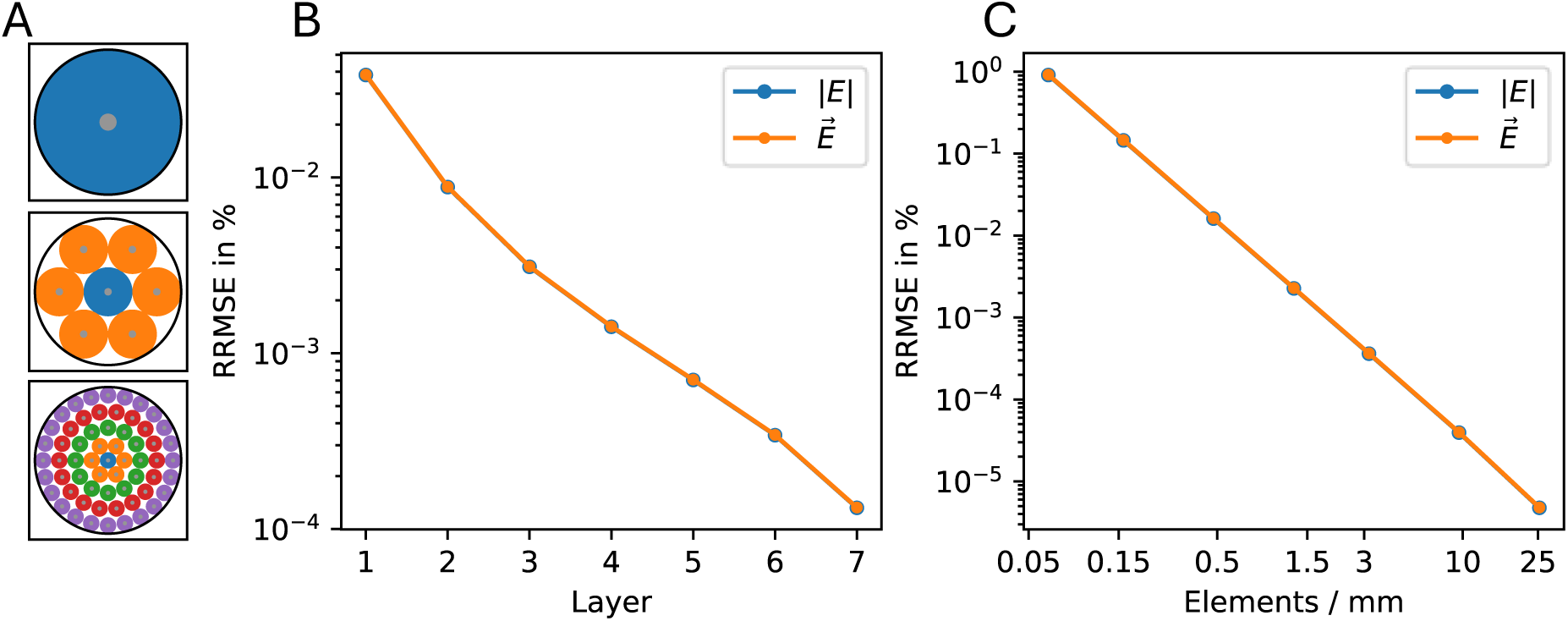
Convergence behavior of the error for the line approximation of a circular coil. A) Layer approximation pattern for 1 layer (top), 2 layers (middle) and 5 layers (bottom). B) Error convergence for an increasing number of layers using the relative root mean square error (RRMSE). Number of lines representing each layer: 1: 1, 2: 7, 3: 19, 4: 37, 5: 61, 6: 91, 7: 127. C) Error convergence for an increasing number of elements per mm.

## Supplementary Material B: Validation of deformation modeling

We approximate coil deformations by concatenated affine transformations, resulting in a low number of free parameters that can be efficiently optimized. To validate this approach, the physical BrainsWay H1 coil was CT-scanned in a deformed state positioned on a football with a 200 mm diameter, and a ground truth model of the deformed coil was generated from the CT scan. The standard H1 coil model was aligned and deformed to match the ground truth model as closely as possible by minimizing the distance between the two coil wires using the DIRECT optimizer implemented in scipy. The electric fields were then simulated on a 200 mm diameter sphere model, corresponding to a version of the simnibs m2m_sphere model scaled to match the size of the football with a diameter of 200 mm. The sphere model had 5 tissue types mimicking the conductivity of human tissues (scalp: 200 mm, 0.465 S/m; bone: 187 mm, 0.01 S/m; CSF: 175 mm, 1.654 S/m; GM 168 mm, 0.275 S/m; and WM 157 mm, 0.126 S/m). The pairwise differences in the electric field magnitudes on the GM surface nodes between the ground-truth H1 model and the deformed and non-deformed H1 models were assessed. The error in the electric field magnitudes between the optimized deformed coil model and ground-truth coil model was reduced to be <2.5%, demonstrating that the approximated deformations represented the true deformed coil shape reasonably well. Supplementary Figure S2 illustrates the differences in coil wires between the initial, ground truth and optimized coil settings (Figure S2A and S2B), and the electric field differences between the coil settings (Figure S2C&D).

**Supplementary Figure S2.**
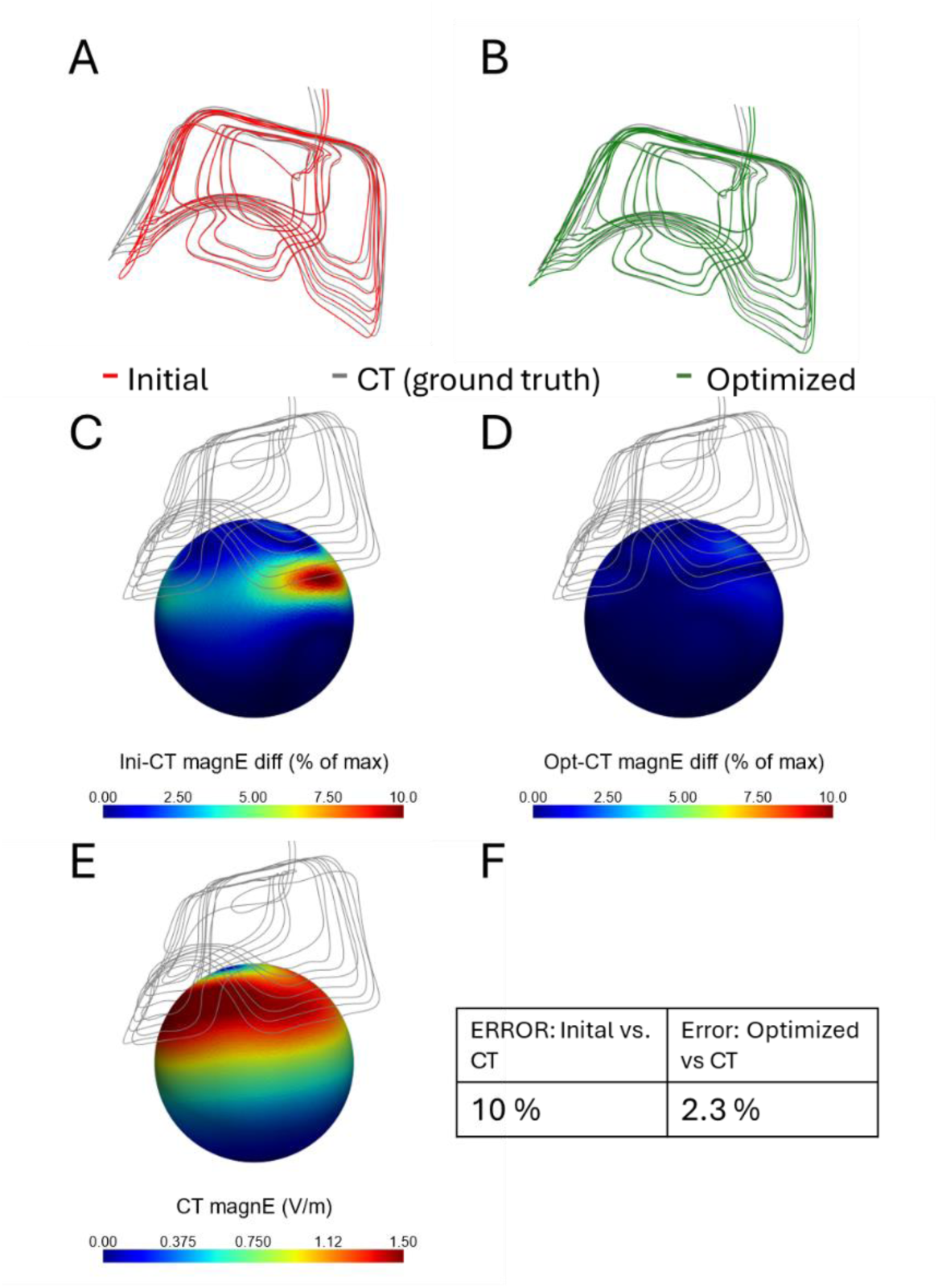
Validation of the deformable computational model of the H1 coil. B) Initial undeformed wire paths (red) and the ground truth (gray) derived from a CT scan in its deformed state on the ball. B) Optimized wire paths (green) and the ground truth (gray). C) Relative difference between the initial coil model and the ground truth. D) Relative difference between the coil model after position and deformation optimization and the ground truth. E) Electric field of the ground truth. F) Average relative differences between the electric fields of the initial undeformed coil model and the ground truth model, as well as of the optimized coil model and the ground truth model.

## Supplementary Material C: Technical notes on the TMS Coil Definition File Format (.tcd)

The TMS Coil Definition File Format (.tcd) is a JSON-based format that defines a single TMS coil. The coil file can be loaded into SimNIBS and used to calculate the magnetic vector potential (A-field) for a given rate of change of the coil current and the magnetic field (B-field) for a given coil current. The following text gives an overview of the format. Details are provided in the SimNIBS documentation as well as in the example scripts for custom coil creation.

**Supplementary Figure S3.**
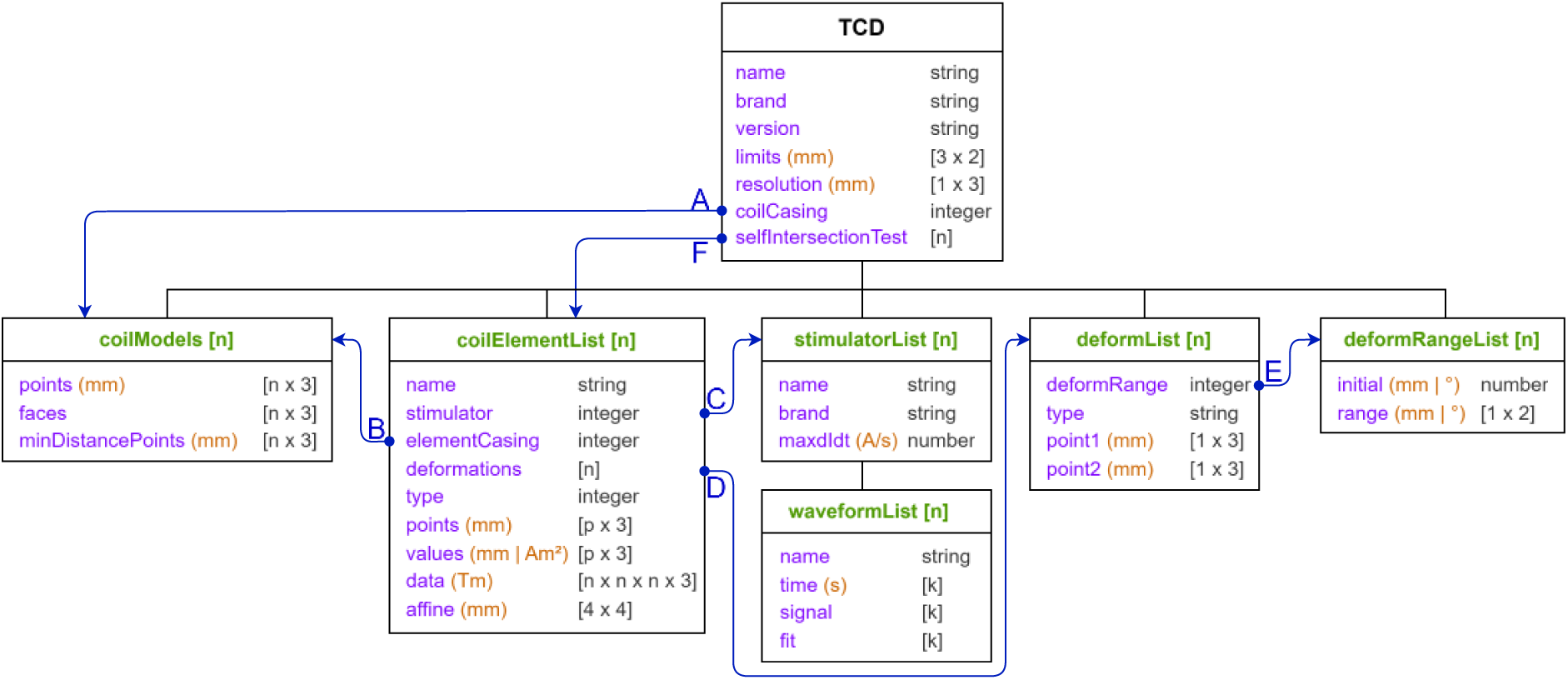
The TMS Coil Definition File Format. Each sub-list (green) can have an arbitrary number of items. A) Each coil can have a general coil casing (or “coilModel”) not associated with any coil element. B) Each coil element can have a casing. C) Each coil element must be connected to one stimulator. D) Each coil element can have an arbitrary number of deformations including no deformations. E) Each deformation uses a deformation range that defines the valid values of the deformation. F) Each coil can have an arbitrary number of self-intersection groups.

The file format holds basic information about the coil, the name, the brand, and the version of the coil file. Its main part is a list of stimulating coil elements (“coilElementList”) that is linked to a list of deformations ("deformList") and a list of geometric models of the sub-casing (“coilModels”). These three components allow for definitions of static coils and coils with one or multiple movable stimulating elements. The following list gives a brief overview of the main components:

- **coilElementList** Each stimulating coil element can use one of three types of stimulating elements:
  - Magnetic dipoles, which are described by their location (in mm), direction, and magnitude (in Am^2^).
  - Line current elements, which are defined by their starting position (in mm), direction, and magnitude (in mm).
  - Magnetic vector potential field (A-field), sampled on a grid, described by the A-field value at each grid location (in Tm) and an affine transformation matrix (in mm).
  Each stimulating coil element has a name, is connected to one stimulator (from “stimulatorList”) (arrow “C” in Suppl. Figure S3), has a casing (from “coilModels”; Suppl. Figure S3 arrow B), and is associated with a list of deformations (from “deformList”) that are applied to this coil element (Suppl. Figure S3 arrow D). The points, values and data lists are stored as plain text or as Base64 encoded and compressed binary. The deformations are applied in the order that they are stored in. Multiple stimulating coil elements can share the same deformation, enabling the common rotation of several coil elements around shared axes.
- **stimulatorList** A list of TMS stimulators is stored to connect different stimulating coil elements to the same stimulator, which can be used in multi-stimulator coil settings. The TMS stimulators are described by their name, brand, maximum dI/dt (in A/s), and a list of waveforms.
- **deformList** A deformation is either defined as a translation in the x, y, or z direction (in mm) or as a rotation around an axis (in degrees) defined by two points (in mm). Its value is defined and bound by a deformation range (Suppl. Figure S3 arrow E).
- **deformRangeList** The limits for deformations are defined as deformation ranges. Multiple deformations, for example, rotations around different axes, can share the same deformation range. These ranges are defined by a minimum and maximum value, and an initial value between the minimum and the maximum.
- **coilModels** For visualization and optimization purposes, the file format contains a list of coil casing triangulated surfaces. Each coil casing surface has a point list (in mm) and a list of faces which are indices into the point list. Additionally, a list of points ("minDistancePoints" in mm) representing the parts of the coil that are supposed to be close to the head can be stored. These are then used for evaluation of the coil-skin distance costs during optimization (Supplementary Figure S4 shows the “minDistancePoints” for the 4 implemented coils as examples). The point and faces lists are stored as plain text or as Base64 encoded and compressed binary.
- **selfIntersectionTest** A list of groups of stimulating coil element indices which will be tested for self-intersections in cases of optimization.

The file format also gives the possibility to store a global coil casing ("coilCasing") that is not associated with a specific stimulating coil element (Suppl. Figure S3 arrow A). In addition, to speed up repeated FEM evaluations for varying coil positions and deformations, the A-fields of the stimulating coil elements can be sampled on 3D grids during preparation. They can then be computationally efficient evaluated for varying coil configurations using simple linear transformations. The file format supports this by giving the option to store information about the axis limits ("limits" in mm) and resolution ("resolution" in mm) of the 3D grids.

**Supplementary Figure S4.**
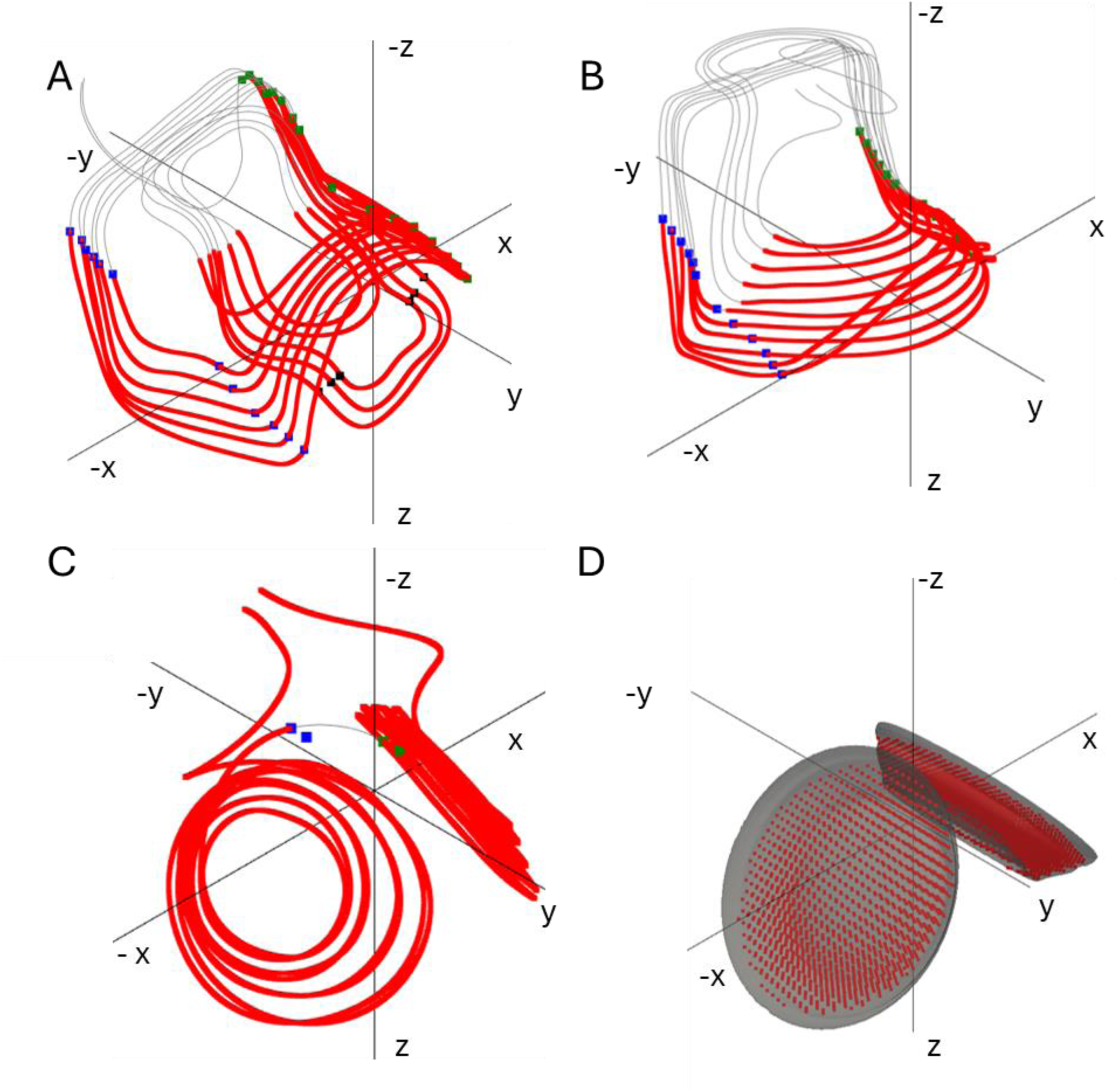
The red **“**minDistancePoints” are used to calculate the *distance* cost during optimization and the points defining the bending axes (blue). The coil wires (H-coils) or coil casing (MST-Twin coil) are shown in gray. The closest distance to the head is determined for each point, and the average then used as *distance* cost (d . A) BrainsWay H1 coil. B) BrainsWay H4 coil. C) BrainsWay H7 coil. D) MagVenture MST-Twin coil.

## Supplementary Material D: Evaluations of the different parts of the cost functions

### Distance Cost

The cost for the distance optimization is the mean distance of pre-selected points on the coil casing or wires (Suppl. Figure S4) to the scalp surface. The points are selected to sample the parts of the coil casing that are facing towards and are close to the scalp. To efficiently calculate the distance, a binary voxel mask of the head is created (resolution 1×1×1 mm³). Using the Euclidean distance transform, the mask is converted into a voxel image where the voxel values describe the signed distance from the scalp surface to the inside and outside of the scalp. The distance at the pre-selected points is cubically interpolated based on the distance voxel image in combination with linear extrapolation.

### E-Field Magnitude Cost

The E-field magnitude cost is the negative of the average E-field strength in a Region of Interest (ROI). To efficiently calculate the E-field magnitude in the ROI, the A-field of each stimulating coil element is sampled on a 3D grid during preparation. The sampled A-field is then used to interpolate the A-field in the head model and update the right hand side of the FEM is to simulate the resulting E-field in the ROI.

### Intersection Penalty

The intersection penalty is the depth-weighted volume intersection between the casings of the coil elements and the head mask. To efficiently calculate the intersection penalty, the scalp surface and the surfaces of the coil element casings are used to create binary voxel masks (resolution 1×1×1 mm³) of the head and casings, respectively. Using the Euclidean distance transform, the head mask is converted into a voxel distance image where outside voxels have a value of 0 and the value of inside voxels represent their distance to the scalp surface. In dependence on the tested coil configuration (i.e., the position relative to the head and the deformations), the transformed coordinates of all voxels of a casing are used to cubically interpolate the depth values from the scalp distance image. This is repeated for all coil elements and the sum of all interpolated depth values is reported as the depth-weighted volume intersection.

### Self-Intersection Penalty

The self-intersection penalty is the volume intersection between selected coil element casings. To efficiently calculate the self-intersection penalty, the surfaces of the coil element casings are used to pre-calculate voxel mask images (resolution 1×1×1 mm³). The coordinates of one coil casing volume are transformed into the space of the other element casing in dependence on the tested coil configuration, and are used to cubically interpolate the volume in another coil casing volume. The sum of these interpolated volume intersections is then used as the volume self-intersection in mm³.

## Supplementary Material E: Additional Figures and Tables

**Supplementary Figure S5.**
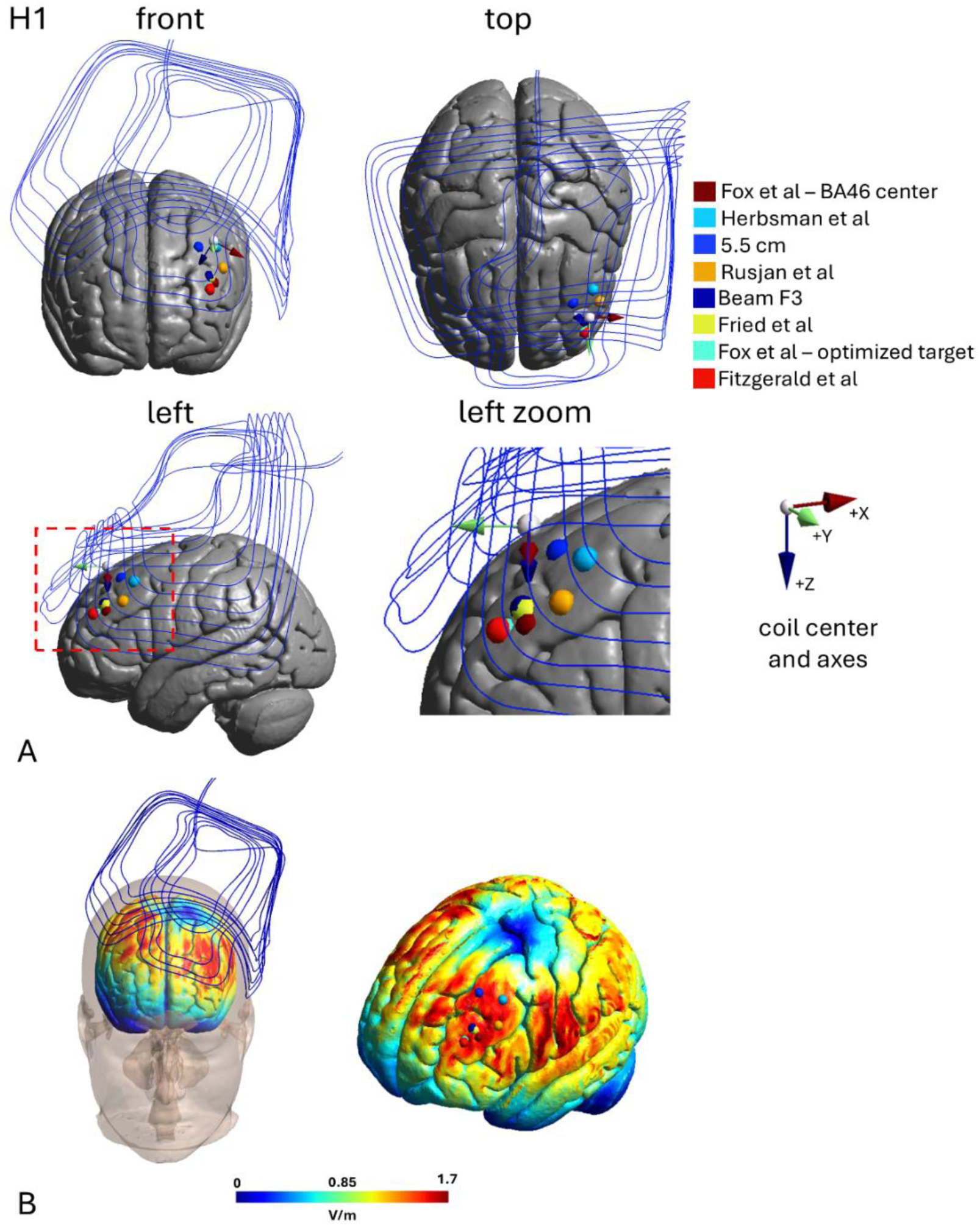
BrainsWay H1 coil on the MNI head. A) Initial coil position. The white sphere indicates the coil center, placed above the center of BA46 (dark red sphere). The visualized target positions were taken from [1,2]. B) Electric field and coil position after distance optimization to resolve intersections of the coil with the head. The coil center was positioned beneath the intersection of the wires running left-to-right and front-to-back. Pilot tests on the MNI head confirmed that the field strength in BA46 decreased when moving the coil center to a more medial position, suggesting that its choice was reasonable. To ensure consistent orientations of the coil axes, they were aligned with the top plate of the coil and the screw threads in the plate (both are not shown here), which are used to secure the coil in the helmet, based on the CT image.

**Supplementary Figure S6.**
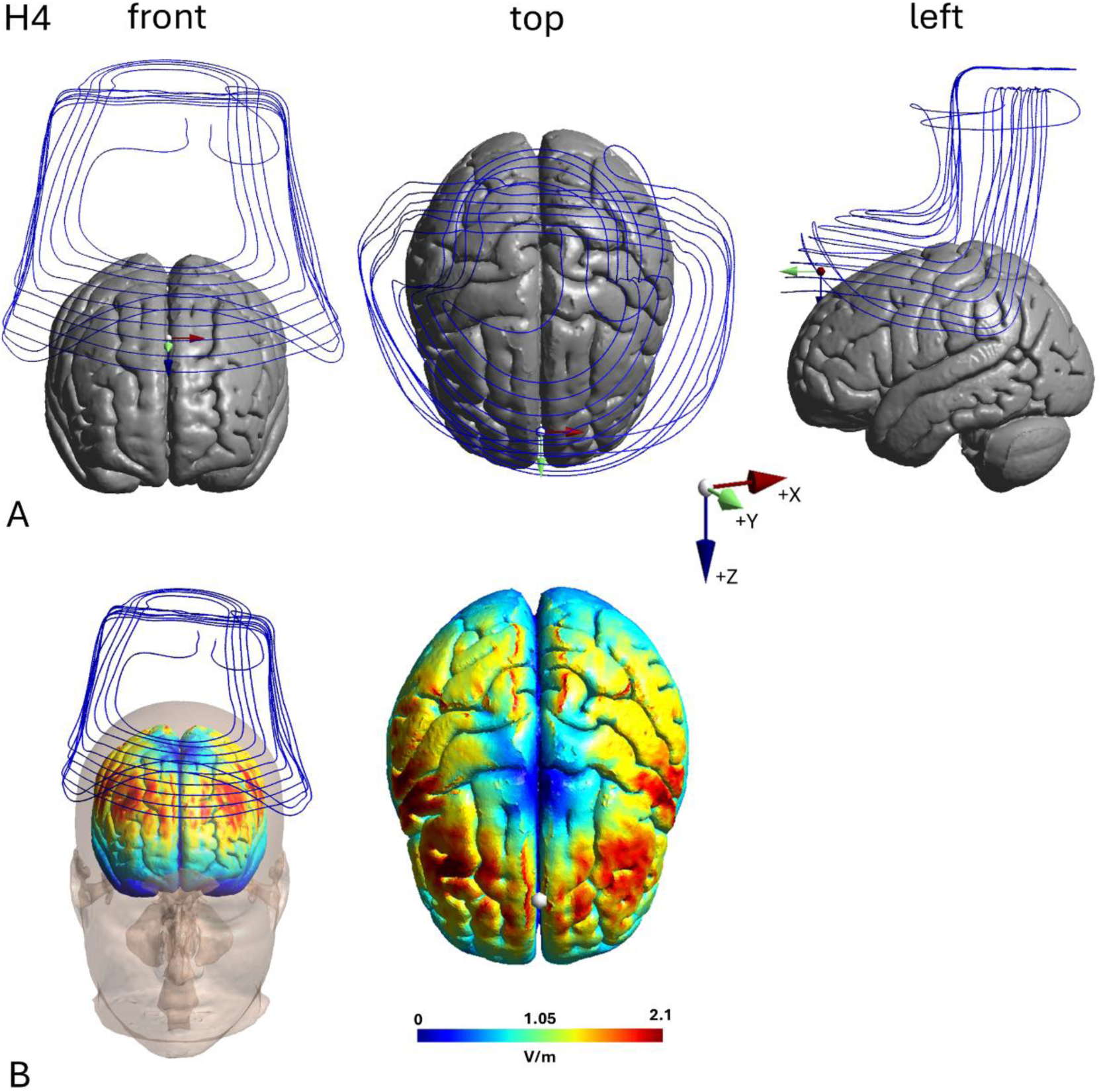
BrainsWay H4 coil on the MNI head. A) Initial coil position. B) Electric field and coil position after distance optimization to resolve intersections of the coil with the head. The white sphere indicates the intended coil center position at MNI coordinate [0, 53, 63] before optimization. The coil axes were aligned with the top plate of the coil.

**Supplementary Figure S7.**
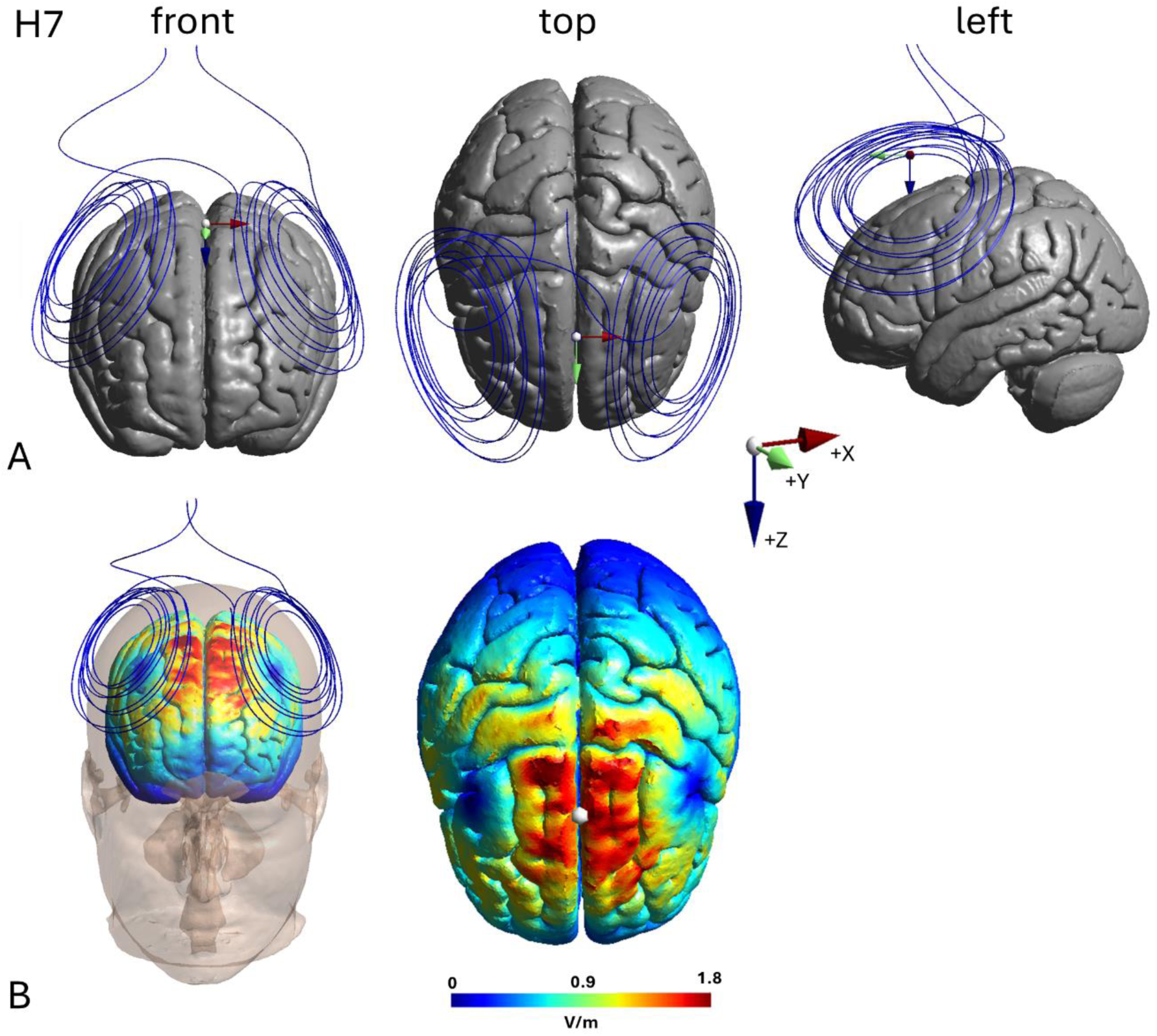
BrainsWay H7 coil on the MNI head. A) Initial coil position. B) Electric field and coil position after distance optimization to resolve intersections of the coil with the head. The white sphere indicates the intended coil center position at MNI coordinate [-5, 20, 87] before optimization. The coil axes were aligned with the top plate of the coil.

**Supplementary Figure S8.**
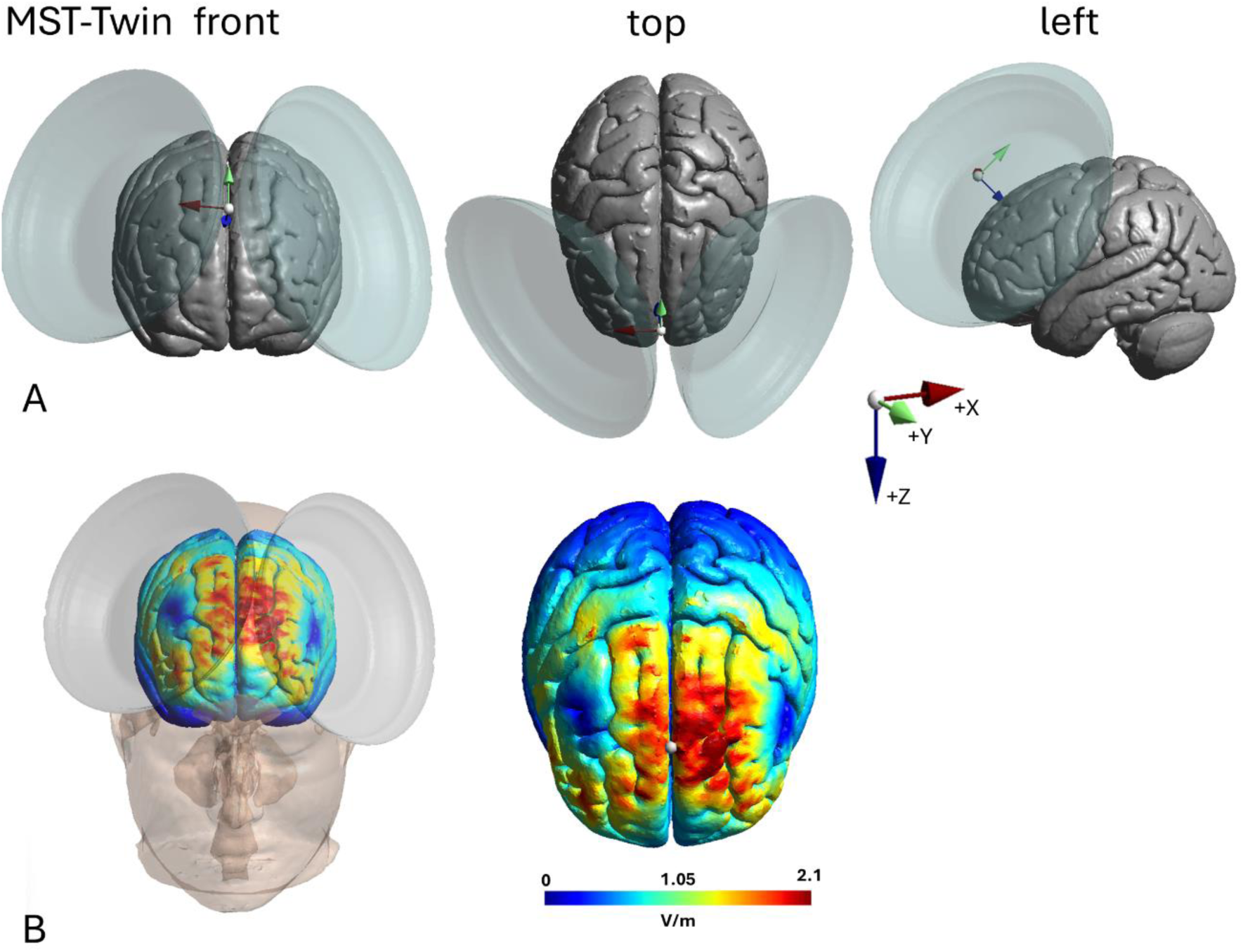
MagVenture MST-Twin coil on the MNI head. A) Initial coil position. B) Electric field and coil position after distance optimization to resolve intersections of the coil with the head. The white sphere indicates the intended coil center position at electrode position Fz before optimization. The coil axes were aligned with the guide rail.

**Supplementary Figure S9.**
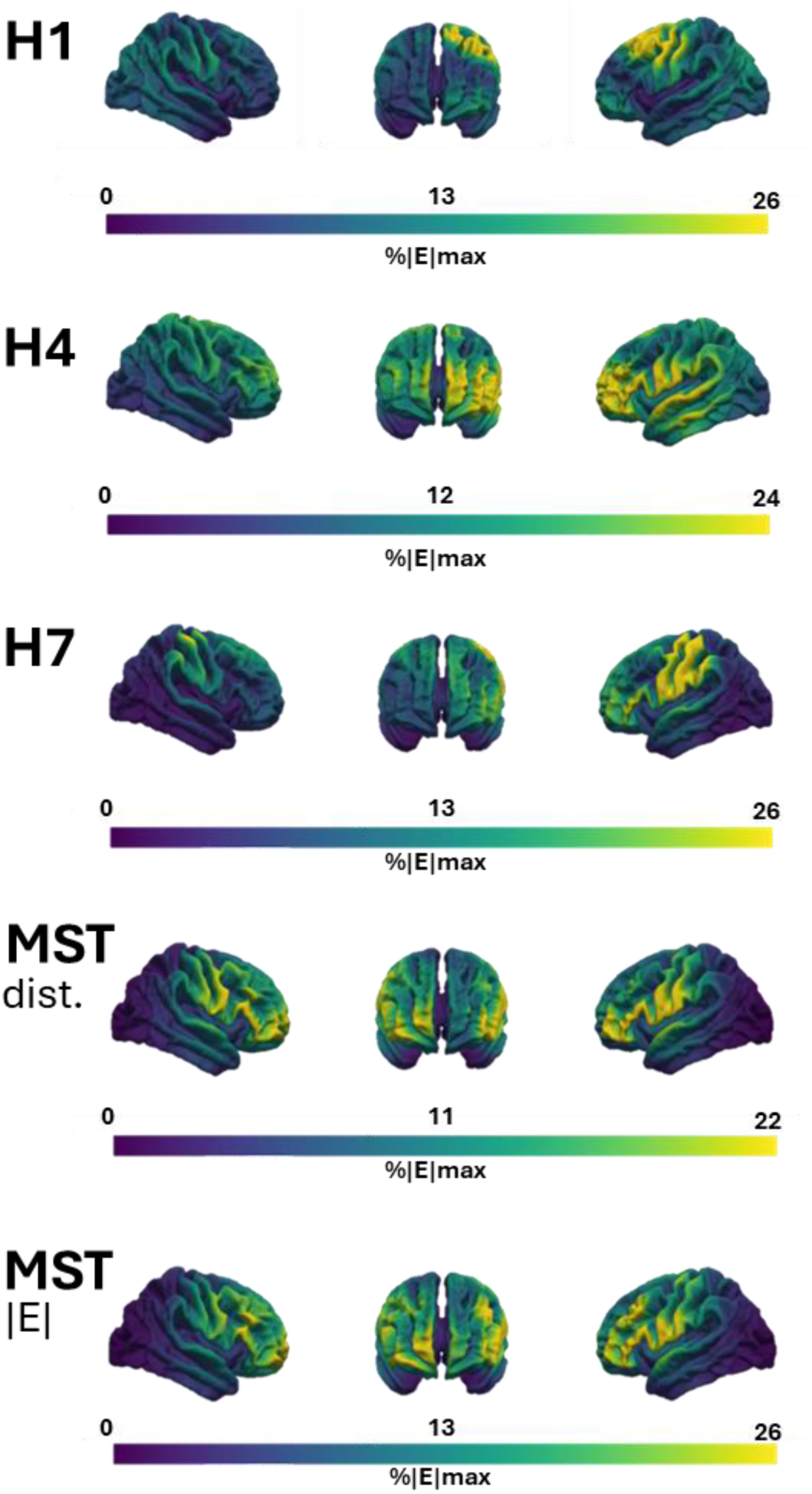
90%iles of the differences between the E-field strength for the initial and optimized coil configurations.

**Supplementary Figure S10.**
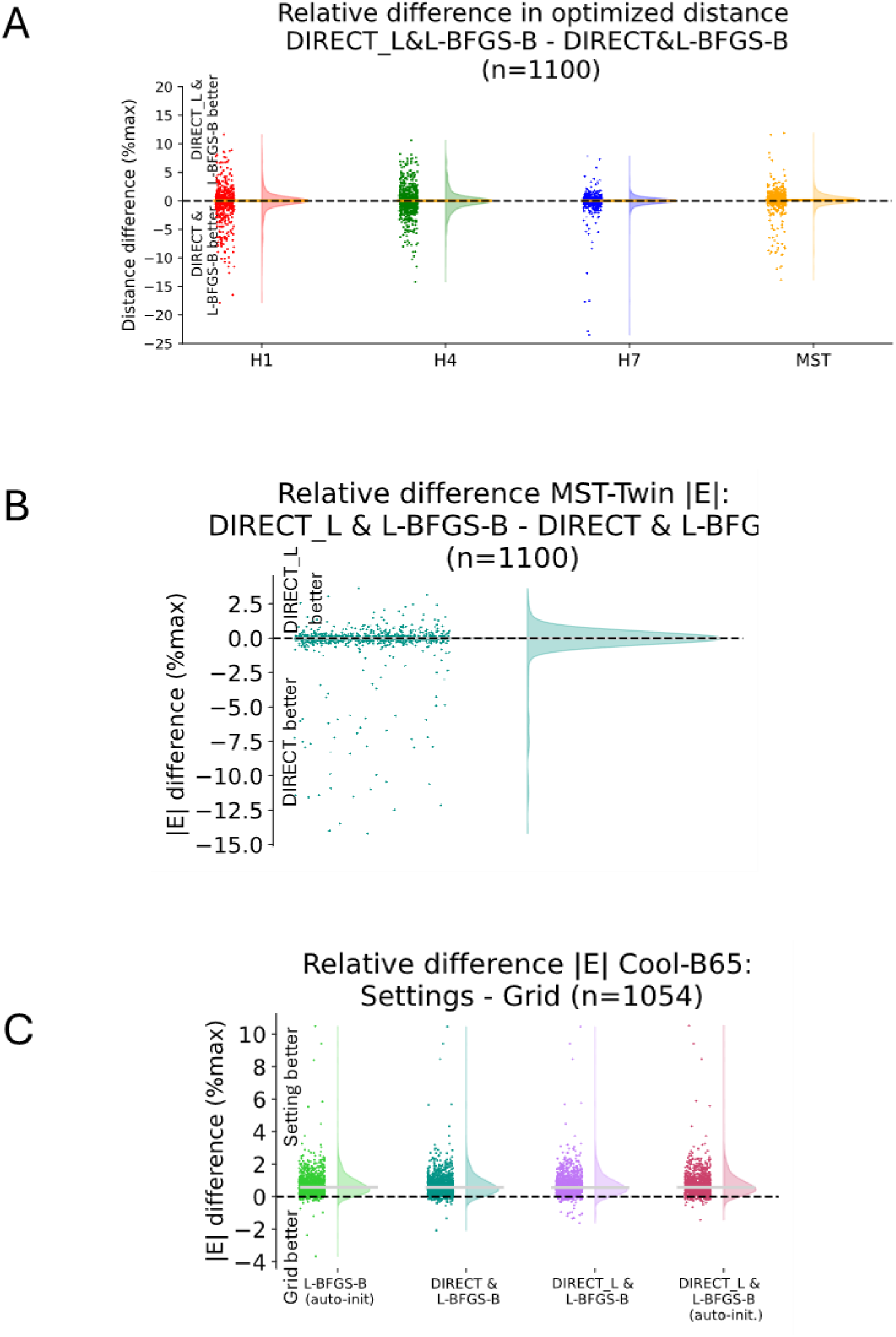
Relative differences between different optimization settings for A) distance optimization, B) E-field optimization of the MST-Twin coil and C) E-field optimization of the Cool-B65 figure-8 coil.

**Supplementary Table S1.** Number of cost function evaluations for the distance and E-field optimization with more exhaustive (DIRECT + L-BFGS-B) and standard (DIRECT-L + L-BFGS-B) search. The results for the settings used in the main paper are marked in **blue**.

| Distance optimization | H1<br>DIRECT +<br>L-BFGS-B | H1<br>DIRECT-L+<br>L-BFGS-B | H4<br>DIRECT +<br>L-BFGS-B | H4<br>DIRECT-L+<br>L-BFGS-B | H7<br>DIRECT +<br>L-BFGS-B | H7<br>DIRECT-L +<br>L-BFGS-B |
| --- | --- | --- | --- | --- | --- | --- |
| N mean ± STD | 7816 ± 3202 | 2137 ± 420 | 6544 ± 2517 | 1479 ± 301 | 5872 ± 1604 | 1423 ± 295 |
| N min | 2957 | 985 | 2436 | 779 | 2095 | 688 |
| N max | 19495 | 3447 | 16804 | 2947 | 12327 | 2990 |

| Distance optimization | MST-Twin<br>DIRECT +<br>L-BFGS-B | MST-Twin<br>DIRECT-L +<br>L-BFGS-B |  | E-field optimization | MST-Twin<br>DIRECT +<br>L-BFGS-B | MST-Twin<br>DIRECT-L +<br>L-BFGS-B |
| --- | --- | --- | --- | --- | --- | --- |
| N mean ± STD | 7213 ± 2263 | 2269 ± 662 | | N mean ± STD | 4921 ± 917 | 2508 ± 722 |
| N min | 3076 | 980 |  | N min | 2056 | 808 |
| N max | 16888 | 4728 |  | N max | 8341 | 5299 |

**Supplementary Table S2.** Number of cost function evaluations for the E-field optimization of the Cool-B65 coil. The results for the settings used in the main paper are marked in **blue**.

|  | L-BFGS-B<br>(auto-init) | L-BFGS-B | DIRECT +<br>L-BFGS-B | DIRECT-L+<br>L-BFGS-B | DIRECT-L+<br>L-BFGS-B<br>(auto-init) | Grid search | High-res<br>grid search |
| --- | --- | --- | --- | --- | --- | --- | --- |
| N mean ± STD | 423 ± 99 | 489 ± 119 | 4750 ± 1207 | 940 ± 169 | 957 ± 151 | 585 ± 0 | 11285 ± 0 |
| N min | 210 | 238 | 2023 | 685 | 719 | - | - |
| N max | 854 | 1015 | 6734 | 2543 | 2213 | - | - |

**Supplementary Table S3.** Timings, number of cost function evaluations and peak memory consumption. Tested on a desktop computer (Ubuntu 22.04, Intel i7-11700, 16 cores, 32GB RAM) using the standard SimNIBS *ernie* head model. The “overhead” covers the time for a final standard FEM simulation with the optimized coil parameters and the creation of visualizations from these simulation results in addition to pre-calculations necessary for estimation of the cost function.

|  | Distance optimization |  |  |  | E-field optimization |  |
| --- | --- | --- | --- | --- | --- | --- |
|  | H1<br>DIRECT-L+<br>L-BFGS-B | H4<br>DIRECT-L+<br>L-BFGS-B | H7<br>DIRECT-L+<br>L-BFGS-B | MST-Twin<br>DIRECT-L+<br>L-BFGS-B | MST-Twin<br>DIRECT-L+<br>L-BFGS-B | Cool-B65<br>L-BFGS-B |
| **Time (optimization)**<br>**in sec** | 68 | 59 | 34 | 43 | 1886 | 338 |
| **Time (overhead)**<br>**in sec** | 83 | 81 | 66 | 64 | 116 | 134 |
| **Total time**<br>**in sec** | 151 | 140 | 100 | 107 | 2002 | 472 |
| **Number of function**<br>**evaluations** | 1390 | 1704 | 1331 | 1359 | 3089 | 686 |
| **Peak memory**<br>**in GB** | 1.7 | 3.6 | 3.5 | 3.8 | 7.5 | 8.2 |

**Supplementary Table S4.**
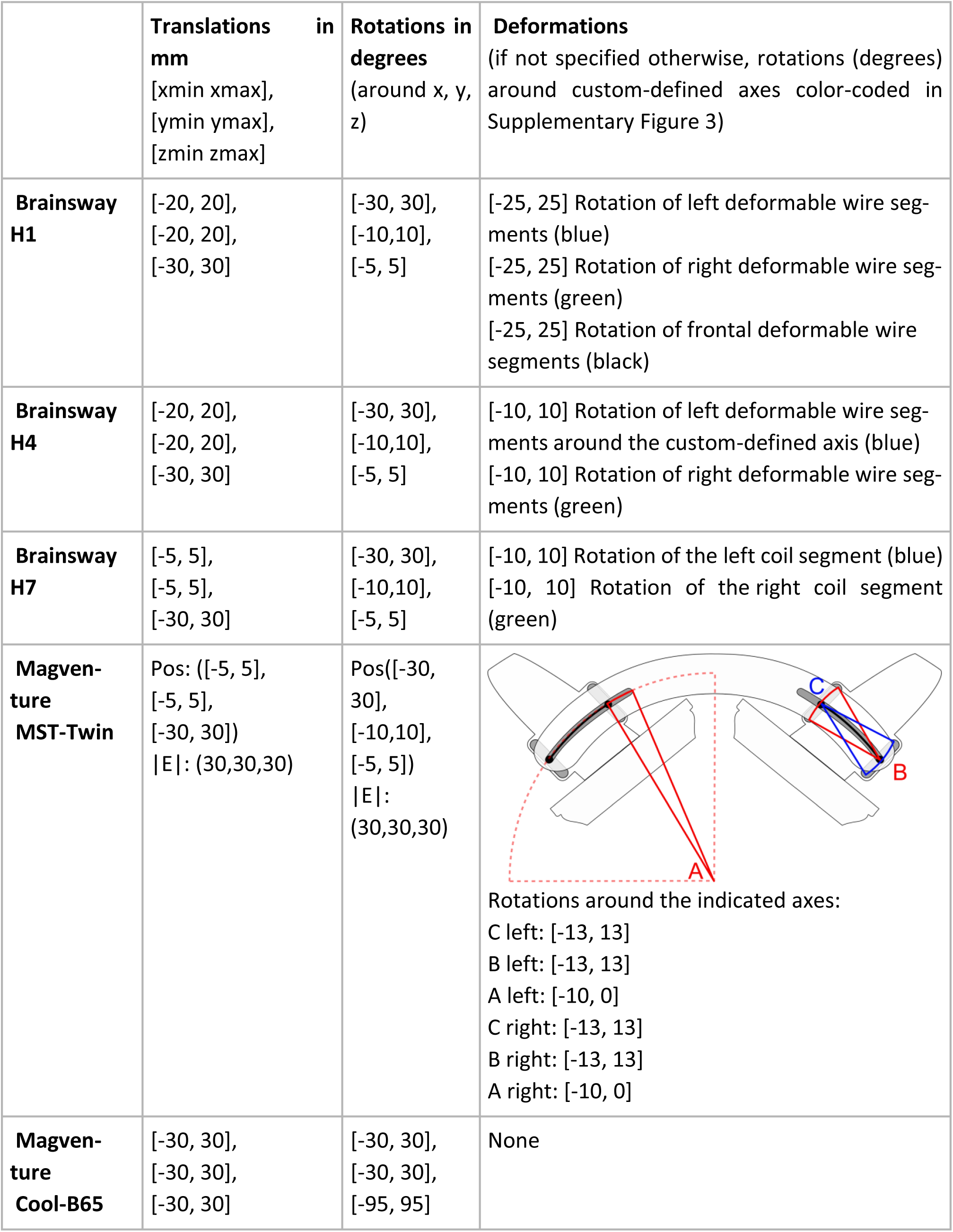
Ranges for the feasible translations and rotations around the coil center and the allowed transformation ranges to mimic the deformation of coil parts.

## Notes

### Competing Interest Statement

The authors have declared no competing interest.

